# First occurrence of *Corynespora cassiicola* infecting chia plant in Bangladesh and its sensitivity to selected fungicides

**DOI:** 10.64898/2026.05.01.722373

**Authors:** Avi Kumar Badhon, Dipali Rani Gupta, Swapan Kumar Paul, Julfikar Ali, Md. Moshiur Rahman, Tofazzal Islam

## Abstract

Chia (*Salvia hispanica* L.) is an emerging crop in Bangladesh valued for its medicinal properties and economic significance. In March 2024, target spot-like symptoms were observed in an experimental chia field (24.75° N, 90.50° E) at Bangladesh Agricultural University in Mymensingh, Bangladesh with disease incidence ranging from 23% to 47% across approximately 0.25 ha. Initially appearing as brick-red spots, these symptoms developed into target-shaped concentric rings, affecting leaves, stems, and inflorescences. A total of 24 fungal isolates were recovered from infected tissue; two representative isolates (BGECh-3 and BGECh-4) were randomly selected for details characterization. Pathogen identity was established through morphological traits, multilocus phylogenetic analysis of internal transcribed spacer (ITS) and elongation factor 1-alpha (*EF-1α*) genes sequence, and pathogenicity confirmation through Koch’s postulates, collectively identifying the causal agent as *Corynespora cassiicola*. The isolates demonstrated a broad host range, successfully infecting brinjal, chili, bottle gourd, country bean, tomato, and soybean. In vitro fungicide sensitivity assays with seven commercial fungicides showed that both isolates were highly sensitive to Goldzim (50% carbendazim), which completely inhibited mycelial growth at 10 µg mL⁻¹. Conza (10% Hexaconazole) and Amister top (18.2% azoxystrobin + 11.4% difenoconazole) reduced growth by up to 85% and 67%, respectively at equal concentration. Other fungicides showed comparatively lower efficacy even at higher concentrations. This study represents the first report of target spot disease of chia caused by *C. cassiicola* in Bangladesh and provides insights for effective disease management strategies.

## Introduction

Chia (*Salvia hispanica* L.), a newly introduced seed crop in Bangladesh, belongs to the family Lamiaceae and has gained considerable attention due to its exceptional nutritional and functional properties [1]. Chia seeds are rich in high-quality fat (30–33%), protein(15–25%), carbohydrates (26–41%), vitamins (A, E, C, B_1_, B_2_, B_3_), antioxidants, and dietary fiber (18–30%), and contain up to 39% oil with an exceptionally high proportion of α-linolenic acid (up to 75%), making chia one of the most efficient plant-based sources of omega-3 fatty acids for food enrichment [2,3]. With the increasing demand for chia as a health-promoting crop, its successful and sustainable cultivation has become an emerging agricultural priority in Bangladesh.

Plant diseases caused by fungal pathogens remain one of the most significant constraints to global crop productivity [4,5]. This challenge is further intensified by climate change, which promotes the geographic expansion of pathogens into previously unsuitable regions and facilitates host shifts to newly introduced crops [6–9]. Such host jumps and long-distance dispersal have been documented in major pathogens, including *Rhizoctonia solani* and *Puccinia graminis* f. sp. *tritici* [10]. Climate change in Bangladesh has significantly impacted crop production and altered disease dynamics by increasing environmental stress on plants and creating favorable conditions for the emergence and spread of plant pathogens. The outbreak of wheat blast in 2016, caused by the fungus *Magnaporthe oryzae* pathotype *Triticum*, is considered a consequence of changing weather patterns and the possible introduction of contaminated seed from abroad [8,11]. Although this pathogen is largely host-specific, which has so far limited its spread to other agriculturally important crops, the situation could become devastating if it were to acquire or exhibit a broader host range affecting crops of major agronomic value.

*Corynespora cassiicola* is a highly polyphagous fungal pathogen reported in more than 70 countries and infecting over 400 plant species, including economically important food, fiber, and horticultural crops [12]. This destructive necrotrophic fungus can infect multiple plant organs, causing characteristic necrotic lesions, premature defoliation, and substantial yield losses [13,14]. The pathogen displays considerable biological and pathogenic variability; isolates often differ markedly in growth rate, sporulation capacity, temperature optima, and virulence, even within similar ecological zones [15,16]. Such variability underscores the importance of accurate pathogen identification and thorough biological characterization, particularly when the pathogen is newly detected on a crop in a specific country. Although *C. cassiicola* has been widely reported worldwide, its occurrence on chia has not yet been documented in Bangladesh. Therefore, a comprehensive understanding of the pathogen associated with target spot of chia in Bangladesh is essential for elucidating disease epidemiology, tracing possible routes of introduction, assessing future outbreak risks, and developing effective management strategies.

The control of fungal diseases relies predominantly on the application of synthetic fungicides. Several broad-spectrum fungicides have been used to manage target spot disease, including succinate dehydrogenase inhibitors (SDHIs) such as boscalid, fluxapyroxad, and fluopyram; quinone outside inhibitors (QoIs) such as azoxystrobin and trifloxystrobin; sterol demethylase inhibitors (DMIs) such as propiconazole, difenoconazole, and tebuconazole; and benzimidazole fungicides such as carbendazim and metalaxyl [17–19]. However, the development of fungicide resistance has been widely documented in diverse *C. cassiicola* populations against many of these active ingredients [17,20,21]. Therefore, understanding pathogen sensitivity to different fungicides and applying appropriate dosages are crucial for effective disease management and minimizing environmental risks.

In this study, typical target spot–infected chia samples were collected during 2024–25 from an experimental field at Bangladesh Agricultural University. The associated fungal isolates were obtained and their pathogenicity was confirmed following Koch’s postulates. Accurate identification of the pathogen was accomplished through detailed morphological characterization combined with multigene phylogenetic analysis. Furthermore, the in vitro sensitivity of selected fungicides was assessed to determine the most effective compounds against the pathogen. The findings of this study provide important insights for developing effective management strategies for target spot disease in Bangladesh.

## Materials and Methods

### Collection and isolation of fungal isolates

The suspected target spot diseased plants were collected from an experimental field of BAU, Mymensingh. The margins of diseased and healthy tissues (leaves, stems, and inflorescences) were cut into 5 mm × 5 mm pieces, followed by surface sterilization with 75% ethanol for 1 minute, rinsing three times with sterile water, and drying on sterile absorbent paper. Infected tissues were placed onto PDA medium and incubated at 25 °C for 2–3 days for fungal growth. Isolates were purified using the single-spore technique on potato dextrose agar at 25 °C in the dark. The isolates were stored at -80 °C for further use.

### Morphological characterization

Morphological characteristics, including colony size, shape, color, and growth rate, were assessed by culturing the isolates on potato dextrose agar (PDA). Briefly, a 5 mm mycelial plug taken from the actively growing margin of a colony was placed at the center of a PDA plate and incubated at 25 °C in the dark for 10 days. Colony diameter was measured in centimeters using a ruler to determine growth rate. Conidia were collected using the whole-mycelium harvest method described by Zhao [22] and filtered through two layers of sterile gauze to remove mycelial debris. The size, shape, and arrangement of conidia and conidiophores were examined under a light microscope (40× magnification) (LEICA DM 1000 Fluorescence microscope), and images were captured using an attached (LEICA MC 190 HD) microscope camera.

### Molecular characterization

Genomic DNA was extracted from two randomly selected fungal isolates (BGECh-3 and BGECh-4) by scraping mycelia from 7-day-old cultures using the Wizard® Genomic DNA Purification Kit (Promega Corporation, Madison, WI, USA), following the manufacturer’s instructions. The internal transcribed spacer (ITS) region of the ribosomal DNA and the translation elongation factor 1-alpha (*EF-1α*) gene were amplified for species-level identification. PCR amplification was performed using the primer pairs ITS1/ITS4 for ITS region [23] and EF595F/EF1160R for (*EF-1α*) gene [24], following established protocols [25]. The primer sequences and expected amplicon sizes are provided in Table 1. Each 50 μl PCR reaction volume containing 5 μl of gDNA (50 ng/μL), 2.5 μl of each forward and reverse primers (10 pmol/μL), 25 μl of 2x GoTaq PCRMasterMix (Promega^TM^ CoTaq C2 green master mix), and 15 μl of nuclease-free water (Promega). For ITS amplification, thermal cycling conditions consisted of an initial denaturation at 95 °C for 2 min; followed by 35 cycles of 95 °C for 40 s, 55 °C for 40 s, and 72 °C for 60 s; and a final extension at 72 °C for 5 min. For *EF-1α* amplification, the program included an initial denaturation at 95 °C for 2 min; followed by 35 cycles of 95 °C for 40 s, 54 °C for 35 s, and 72 °C for 60 s; and a final extension at 72 °C for 5 min. The amplified PCR products were visualized under a UV transilluminator, purified, and sent for sequencing to Apical Scientific Sequencing. The obtained sequences were compared with reference ITS and *EF-1α* sequences of *C. cassiicola* available in the NCBI GenBank database using the BLAST algorithm. Multiple sequence alignment and homology analyses were performed using MAFFT version 7 [26]. The newly generated sequences were deposited in GenBank. Phylogenetic relationships were inferred using the maximum likelihood method implemented in MEGA version 11 with 1,000 bootstrap replicates [27].

**Table 1.**
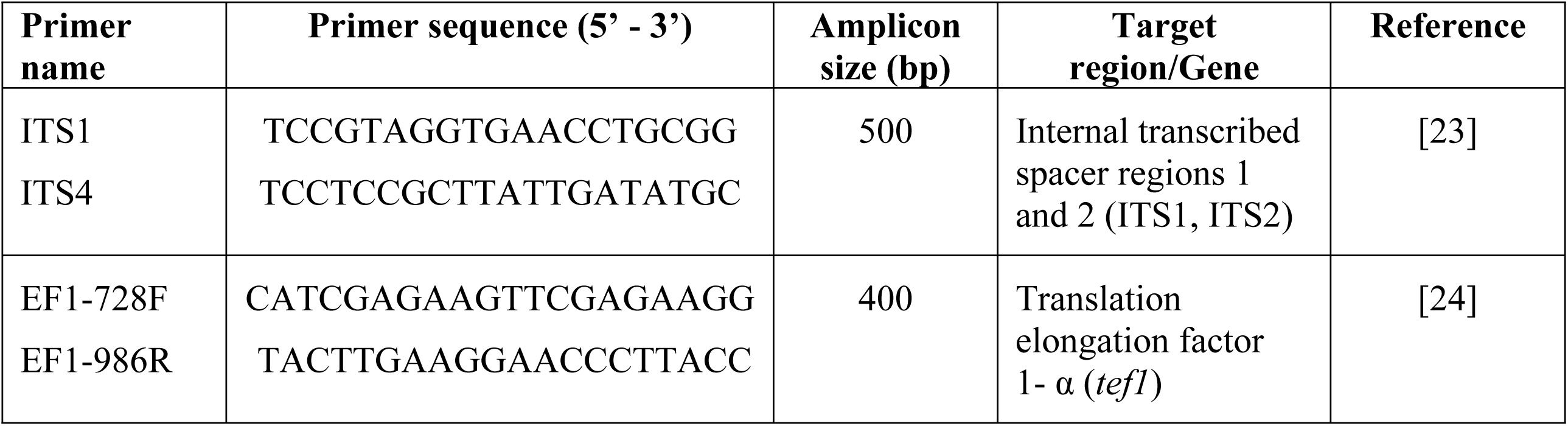
Primer used for PCR amplification and DNA sequencing for molecular identification of fungal isolates.

### Inoculum Preparation and pathogenicity assay

For the preparation of fungal inoculum, the isolates were cultured on potato dextrose agar (PDA) at 25 °C in the dark for 10 days. The mycelial mats were then flooded with 15 mL of sterile distilled water containing 0.1% Tween 20 and gently brushed with a sterile soft brush to release the conidia. The resulting suspension was filtered through two layers of sterile gauze into 50 mL Falcon tubes to remove mycelial debris. The conidial concentration was adjusted to 1 × 10⁴ spores/mL using a hemocytometer prior to inoculation.

For pathogenicity assays, 1 mL of conidial suspension (1 × 10⁴ spores/mL) from each of the two *C. cassiicola* isolates was sprayed onto one-month-old chia seedlings. Each isolate was applied to three independent plants per assay, and the experiment was repeated six times to ensure reproducibility. Additionally, inoculations were performed on stems and inflorescences. Control plants were sprayed with sterile distilled water containing 0.1% Tween 20. After inoculation, plants were maintained in a growth chamber at 25 ± 2 °C under a 16 h light/8 h dark photoperiod and high relative humidity (>90%) for 48 h to facilitate infection. Symptom development was monitored at every two-day interval. To fulfill Koch’s postulates, the pathogen was reisolated from symptomatic tissues of artificially inoculated plants, purified on PDA plates for confirmation.

To assess the host range of *C. cassiicola* isolates BGECh-3 and BGECh-4, were inoculated onto representative vegetable crop commonly cultivated in Bangladesh, including tomato (cv. BARI Tomato-5), chili (cv. KARASHI 5424), brinjal (cv. BARI Begun-6), soybean (cv. BARI Soybean-5), country bean (cv. BARI Seam-1), and bottle gourd (cv. BARI Lau-1). Seedlings were grown in plastic pots containing potting mix, and 20–30 days aged plants were inoculated with the pathogen following the procedure described above. Each treatment included appropriate controls, and symptom development was monitored under controlled conditions.

### Fungicides sensitivity assay

The sensitivities of *C. cassicola* isolates to seven fungicides were assessed by determination of their effective concentration using poisonplate assays as described by [28,29]. The fungicides tested include carbendazim (Goldazim ® 500 SC, Square Pharmaceuticals PLC, Bangladesh), mancozeb (Ticozeb ® 80 WP, Square Pharmaceuticals PLC, Bangladesh), hexaconazole (Conza Plus 10 SC, ACI crop Care), propiconazole (Tilt 250 EC, Syngenta), 50% Tebuconazole combine with 25% Trifloxystrobin (Nativo 75 WG, Bayer Crop Science), combination of 18.2% Azoxystrobin and 11.4% Difenoconazole (Amister Top 325 SC) and 37.5% Carboxin combined with 37.5% Thiram (Provax 200 WP, Hossain Enterprise C.C. Limited).

Stock solution (1000 μg ml^−1^) of each fungicide was prepared in sterile distilled water and incorporated into autoclaved PDA medium after cooling to 50 to 55°C to obtain finial concentrations 0.1, 1, 10, 50 and 100 μg ml^−1^. Fungicide-amended media were poured into Petri dishes, while non-amended PDA served as the control. Subsequently, a 5-mm mycelial plug was taken from actively growing edge of a 7-day-old culture, placed at the center of each plate, and incubated in the dark at 28°C for 10 days.

Colony diameter (minus the original diameter of the inoculation plug) was measured along two perpendicular axes, and mean value was used to calculate radial growth according to [30]. Relative growth inhibition (RGI) was calculated according to the following formula:

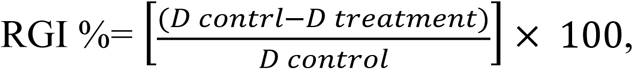

where D = diameter of the colony. Each isolate was tested for 6 replicates and the experiment was repeated twice independently.

The effective concentration to inhibit 50% of mycelial growth (EC_50_) was estimated by fitting a four-parameter log-logistic dose-response model using the ‘drc’ package in R software [31]. EC_50_ values were derived from the fitted curves based on relative growth inhibition data, and model parameters were used to describe the dose-response relationship for each fungicide.

### Statistical analysis

All experiments were conducted using a completely randomized design (CRD) with six replicates per treatment. Data were analyzed using the R software (version 4.4.2). Data on disease incidence, disease severity, and fungicide sensitivity (EC₅₀) were analyzed using analysis of variance (ANOVA) to determine significant differences among treatments. Mean comparisons were performed using Fisher’s least significant difference (LSD) test at *P* ≤ 0.05, and results are presented as mean ± standard error (SE).

## Results

### Disease symptoms and severity

In March 2024, distinctive target spot symptoms were observed in an experimental chia field (24.75° N, 90.50° E) at Bangladesh Agricultural University in Mymensingh, Bangladesh. The disease, initially detected during the vegetative stage, worsened significantly during flowering stage. The initial symptoms manifested as brick-red spots (Fig 1A), these symptoms developed into target-shaped concentric rings, affecting leaves, stems, and inflorescences (Fig 1B,C) indicating the ability of the pathogen to infect multiple aerial parts of the host. Approximately 0.25 hectares of chia field was affected with disease incidence ranging from 23% to 47%. A total of 24 fungal isolates were collected from eight infected chia plant, with at least three from each sample.

**Fig 1.**
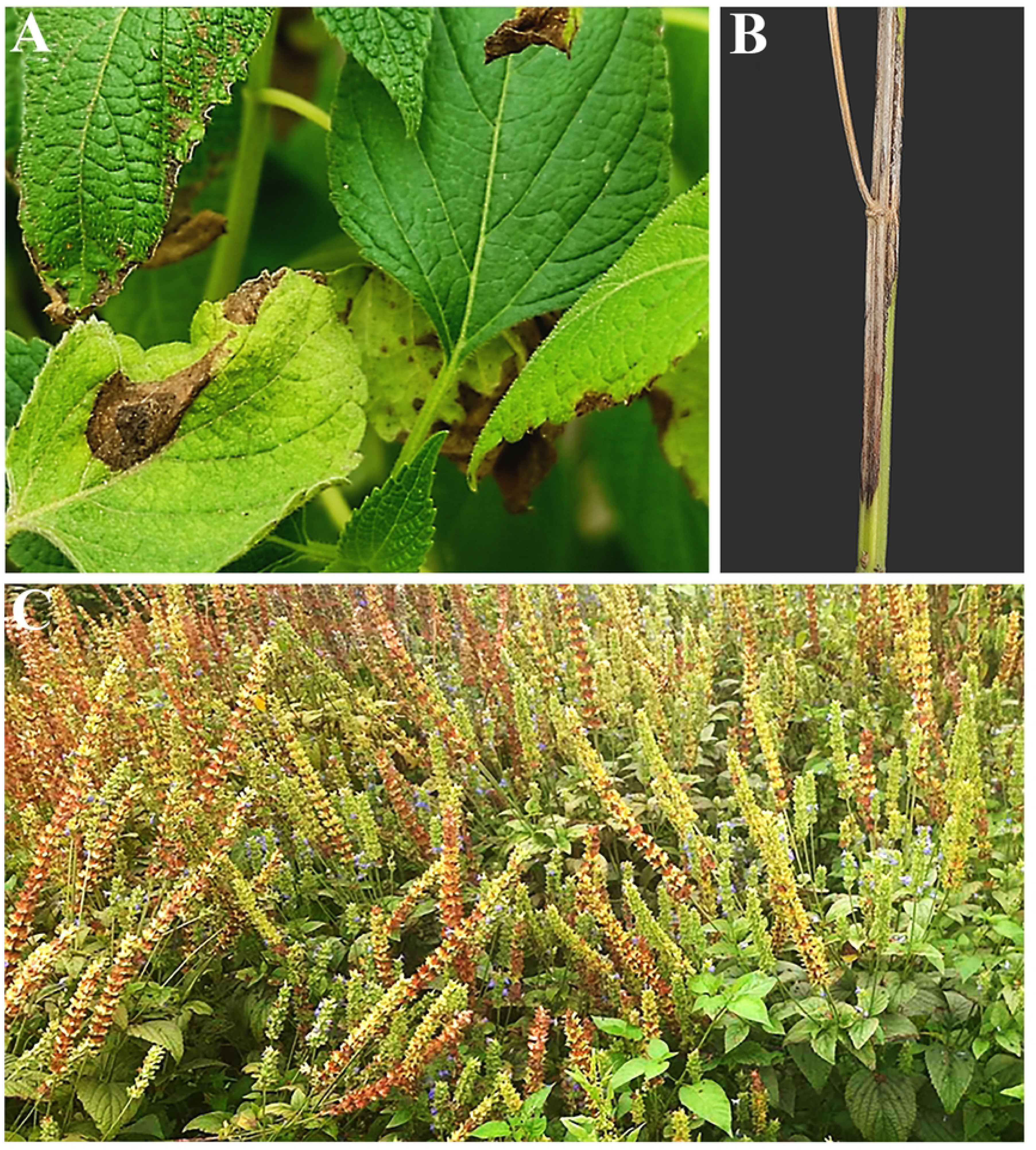
Symptoms of target spot disease caused by *C. cassiicola* on chia under field conditions. (A) necrotic, circular to irregular lesions with concentric rings on leaves; (B) elongated dark lesion on stems; and (C) infected inflorescences showing blight symptoms in a naturally infected field.

### Morphological characterization

Fungal colonies on potato dextrose agar (PDA) were initially olivaceous, gradually turning brown with age, and exhibited an average radial growth rate of 0.8 cm per day (Fig 2A,B). The conidiophores were straight to slightly curved, smooth, unbranched, and grayish-brown in color (Fig 2C). No significant variation in conidial size or shape was observed among the isolates examined. Conidia were produced either singly or in chains on the conidiophores (Fig 2C,D). Although different conidial shapes were observed, straight forms predominated over obclavate or cylindrical types (Fig 2E). The conidia were obclavate to cylindrical in shape and contained 2 to 10 pseudosepta, most commonly 2–5. Measurements of 100 conidia revealed a size range of 9–130 μm in length and 5–12 μm in width (Fig 2D–G). Based on these morphological characteristics, the isolates were identified as *Corynespora* sp [32].

**Fig 2.**
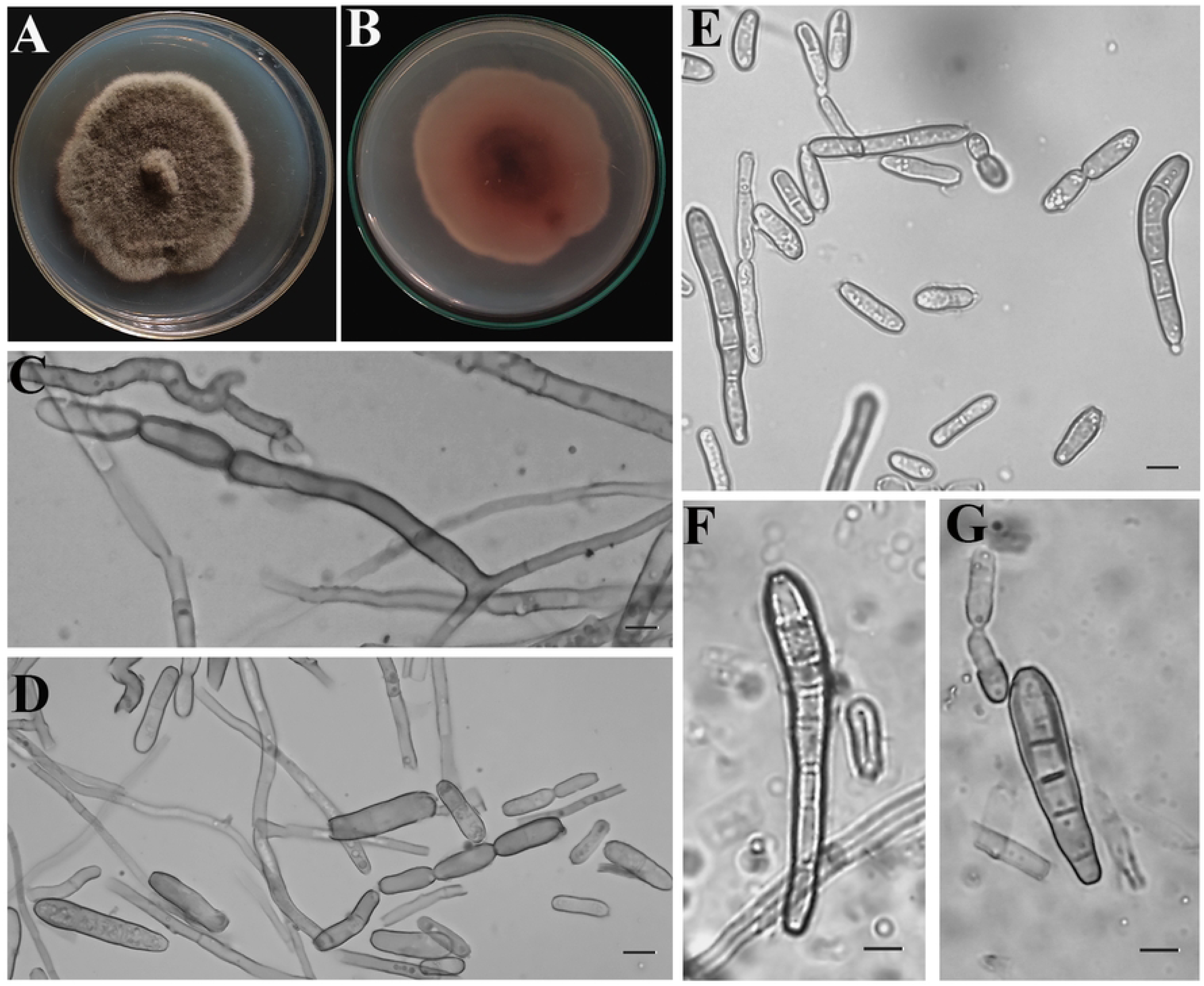
Morphological characteristics of *C. cassicola* isolates BGECh-3 culture on PDA medium collected from diseased chia plant. (A) colony morphology (observed view); (B) reverse view; (C) conidia with conidiophore; (D) chain-liked conidial arrangement; (E) conidia with various size and shape; (F–G) obclavate-shaped conidia.

### Molecular identification of fungal isolates

The internal transcribed spacer (ITS) region and the translation elongation factor 1-alpha (*TEF 1-α*) gene are widely recognized as reliable DNA barcodes for fungal species identification [25,33]. To confirm the identity of the isolates, both ITS and *TEF 1-α* regions were amplified, sequenced, and compared with reference sequences available in the NCBI GenBank database. The newly generated sequences were deposited in GenBank and assigned accession numbers PQ588139 and PQ588449 for the ITS region, and PQ998226 and PQ998227 for the *TEF 1-α* gene. BLASTn analysis showed that the ITS sequences shared 100% identity with *C. cassiicola* isolate CC-01 (KP759967), while the *TEF 1-α* sequences exhibited 99.69% similarity with isolate Cc_01 (MK882240). Furthermore, maximum likelihood phylogenetic analysis based on concatenated ITS and *TEF 1-α* sequences demonstrated that isolates BGECh-3 and BGECh-4 clustered with reference strains of *C. cassiicola*, thereby confirming their taxonomic identity (Fig 3).

**Fig 3.**
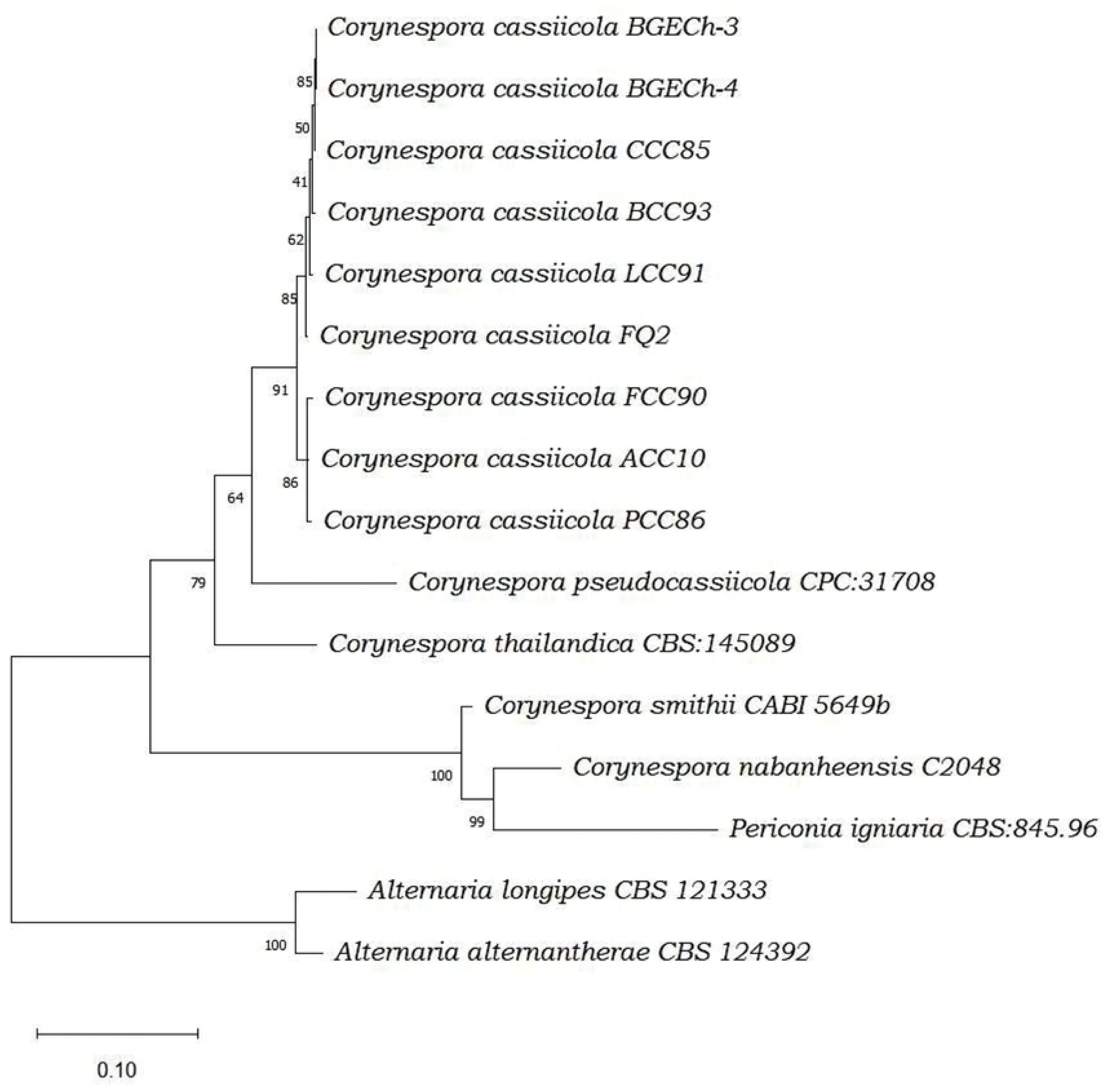
Phylogenetic tree of *C. cassiicola* using combined ITS-rDNA and *TEF1-α* gene sequences by Maximum likelihood method. The tree showed that the sequence of present isolates BGECh-3 and BGECh-4 shared a common clade with reference sequence of *C. cassicola*. Sequences from the present study are inducted by colored block. The percentage of replicate trees in which the associated taxa clustered together in the bootstrap test (1000 replicates) are shown below the branches. The tree is drawn to scale, with branch lengths measured in the number of substitutions per site. This analysis involved 16 nucleotide sequences. There were a total of 1060 positions in the final dataset.

### Pathogenicity assay

Inoculated seedlings developed typical target spot symptoms within 6–8 days, and inflorescences and stems within 10–15 days (Fig 4B,C), while controls remained symptomless (Fig 4A). The pathogen was successfully re-isolated from symptomatic tissues of all inoculated plants (Fig 4D,E) confirming *C. cassiicola* as the causal agent of target spot of chia by morphological and molecular characterization as described above. However, no *C. cassiicola* was recovered from healthy plants. These results indicate that the isolates BGECh-3 and BGECh-4 are the causal agent of target spot disease in chia in Bangladesh.

**Fig 4.**
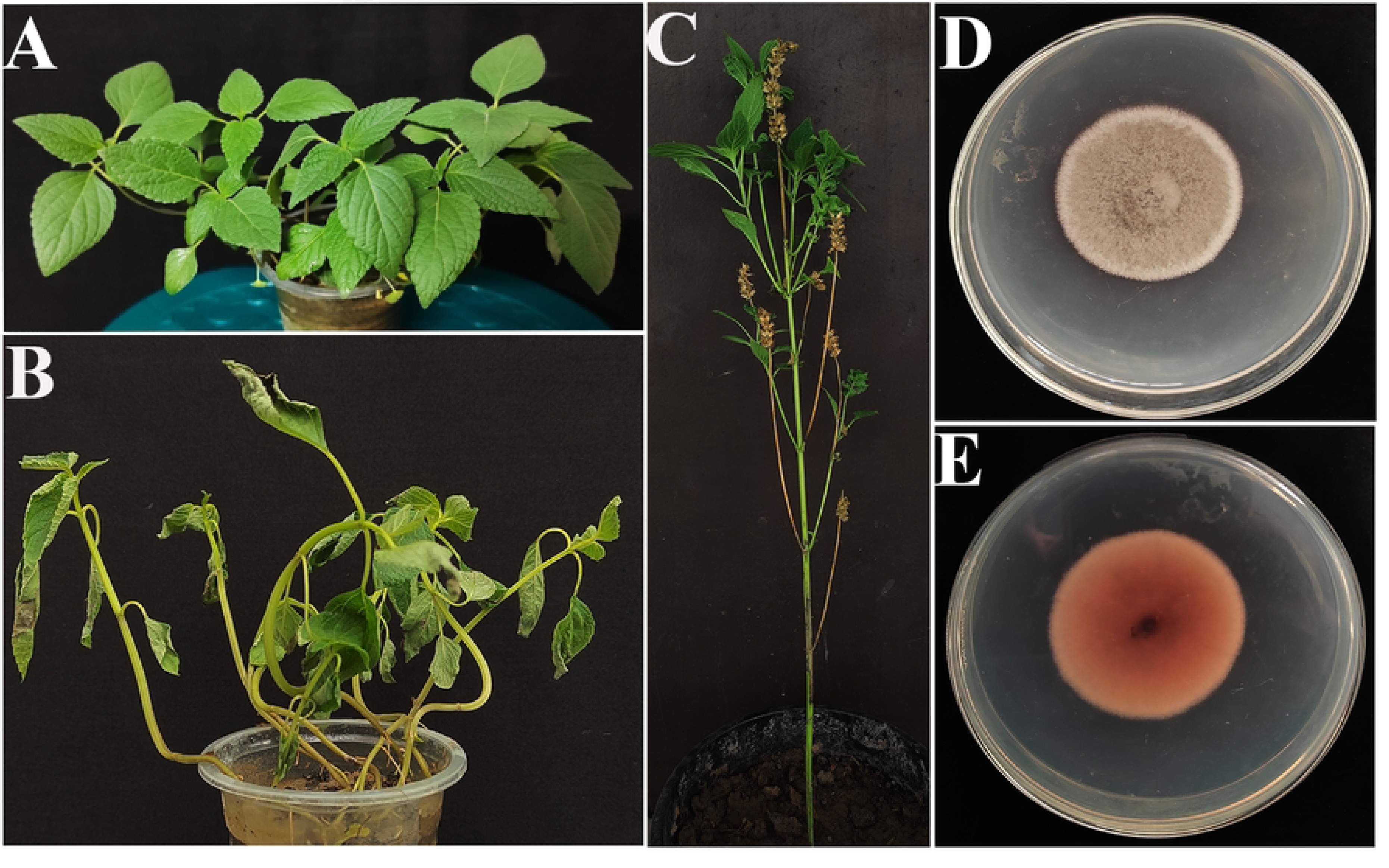
Disease symptoms on chia plant inoculated with *C. cassiicola* isolate BGECh-3 under controlled conditions (10-15 dpi). (A) water treated control plants remained symptomless; () inoculated seedlings showing typical disease symptoms; (C) infected inflorescence; (D-E) colony morphology of reisolated pathogen, recovered from the artificially inoculated plant (observed and reverse views).

### Host range of *C. cassicola* isolates

Both the isolates were found to be highly pathogenic on chili, tomatoes, country beans, sweet gourd, brinjal and soybean under inoculated condition. Isolates produced typical symptoms on the leaves of the inoculated plants. In Tomato, eggplant and bottle gourd, severe disease symptoms were observed, and the plants became collapsed within 10 days after pathogen inoculation (Fig 5A,C,E). In soyabean, initial symptoms appeared 5 days after inoculation, beginning as brown discoloration on older leaves. As the disease progressed, whole plants gradually turned yellow and eventually dried (Fig 5G). In contrast, symptom development was slower in chili and country bean, where characteristic target spot symptom with concentric rings developed approximately 12-14 days after inoculation (Fig 5I,K).

**Fig 5.**
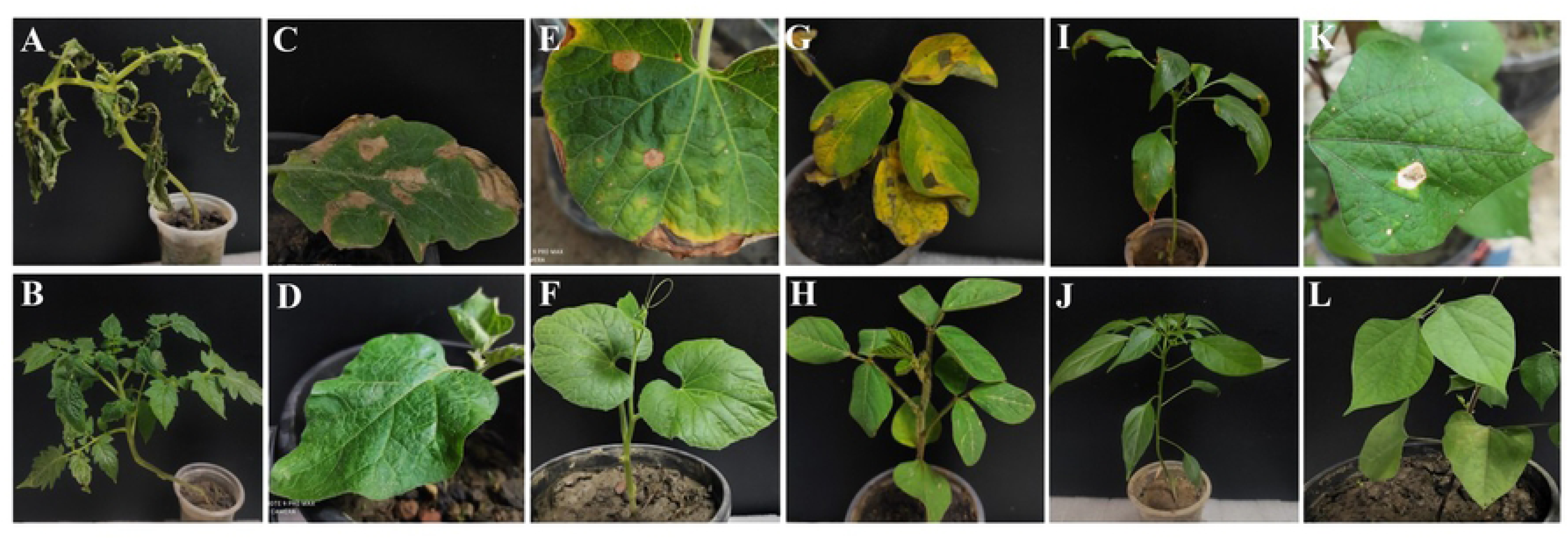
Pathogenicity assessment of *C. cassiicola* on multiple host plants under control condition. Respective symptoms of target leaf spots are presented on inoculated plant (A) tomato, (C) brinjal, (E) bottle-gourd, (G) soyabean, (I) chili, (K) country-bean. Corresponding non-inoculated control plants are presented directly below each treatment: (B), (D), (F), (H), (J), and (L), respectively. Symptom development was monitored at 3-day intervals. Typical necrotic lesions with concentric rings were observed in all inoculated plants, whereas no symptoms developed in the control plants.

### Fungicide sensitivity assays

Fungicide sensitivity was evaluated based on mycelial growth inhibition. Both *C. cassiicola* isolates showed similar sensitivity to the tested fungicides, and the data were then pooled. A clear dose-dependent inhibition effect was observed for most of the fungicides, whereas Ticozeb (mancozeb), showed minimal inhibitory effect on the mycelial growth across the tested concentrations. Among the fungicides, the systemic fungicides Goldazim (carbendazim) and Conza (Hexaconazole) exhibit the higher efficacy. At a concentration of 10 μg mL⁻¹, both fungicides produced significantly higher inhibition percentages compared with the other treatments. At this concentration, Conza inhibited mycelial growth by 85–88%, whereas Goldazim achieved complete (100%) inhibition (Fig 6). Amistar Top (azoxystrobin + difenoconazole) and Tilt (Propiconazole) produced moderate inhibition (67-69% and 58-61%, respectively) at the same concentration (Fig 6) whereas, Nativo (Tebuconazole + Trifloxystrobin) and Provax (Carboxin + Thiram) yielded lower levels of growth inhibition (approximately 50% and 45-47%, respectively) at 10 μg mL⁻¹ (data not shown).

**Fig 6.**
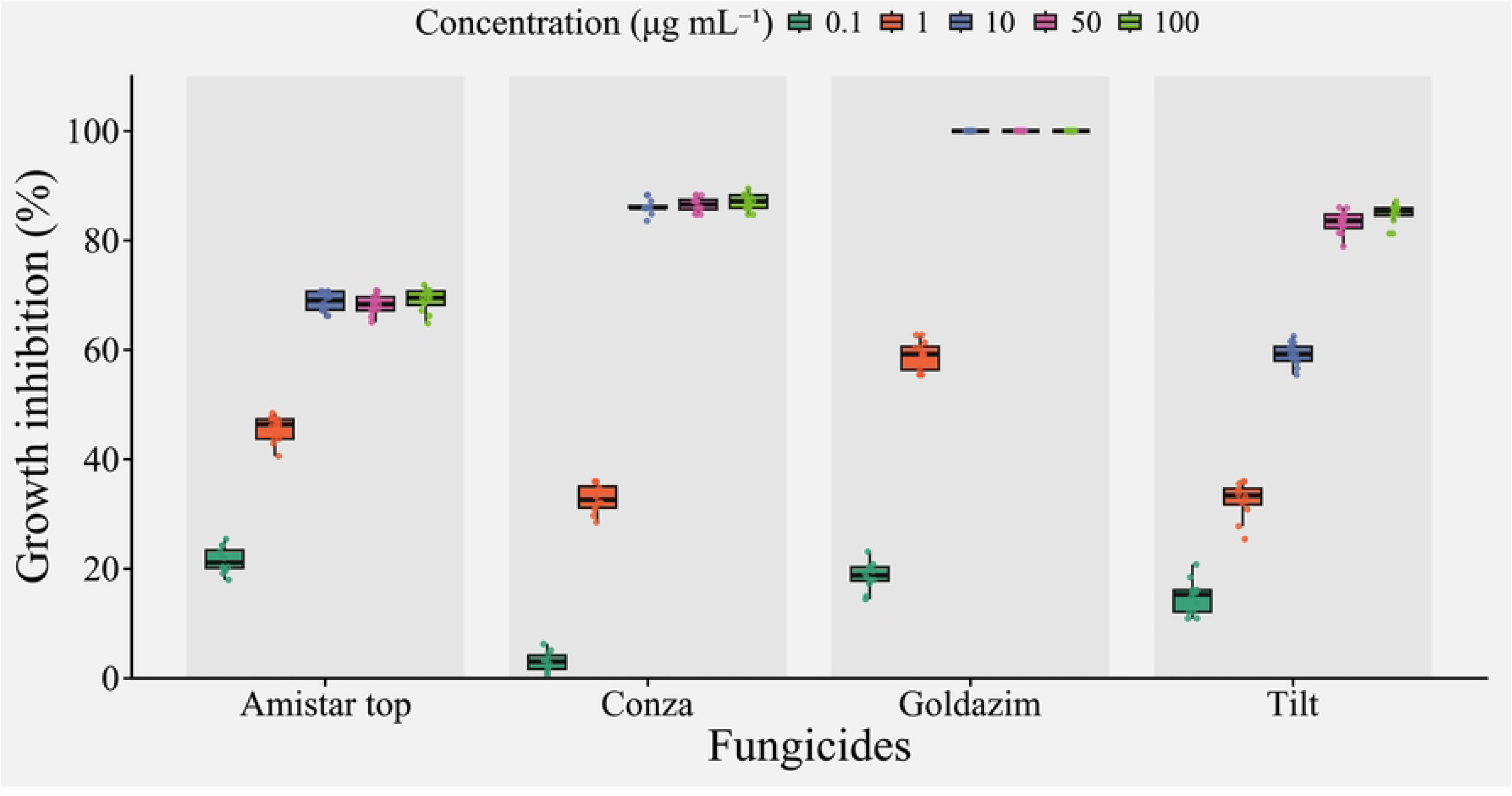
Mycelial growth inhibition (%) of *C. cassiicola* under four fungicides across five concentrations (0.1, 1, 10, 50, and 100 μg mL⁻¹). Boxplots represent the distribution of inhibition values for Amistar Top, Conza, Goldazim, and Tilt based on six independent replicates per treatment, with overlaid points indicating individual observations. The central line denotes the median, boxes indicate the interquartile range (IQR), and whiskers denote the range of observed values.

Notable, increasing the concentration of Conza and Amistar Top did not result in proportional increase in mycelial growth inhibition, indicating a plateau in efficacy under the tested conditions. On the other hand, Ticozeb exhibited the weakest antifungal activity, with limited inhibition at the concentration 10 μg mL⁻¹ and increasing the concentration upto 100 μg mL⁻¹ resulted in 70% reduction in mycelial growth (data not shown).

Similar trends were observed in the dose-response analysis (Fig 7). The fitted models indicated concentration-dependent inhibition for all fungicides, with clear differences in EC_50_ values. Goldazim exhibited the highest activity, with EC₅₀ values of 0.76 μg mL⁻¹, followed by Amistar Top (1.07 μg mL⁻¹) and Conza (1.41 μg mL⁻¹). In contrast, Tilt showed comparatively lower activity, with a higher EC₅₀ value of 4.45 μg mL⁻¹. These results confirm that carbendazim- and hexaconazole-based fungicides are more effective in inhibiting *C. cassiicola* than propiconazole under the tested conditions and further indicate variability in fungicide sensitivity among treatments.

**Fig 7.**
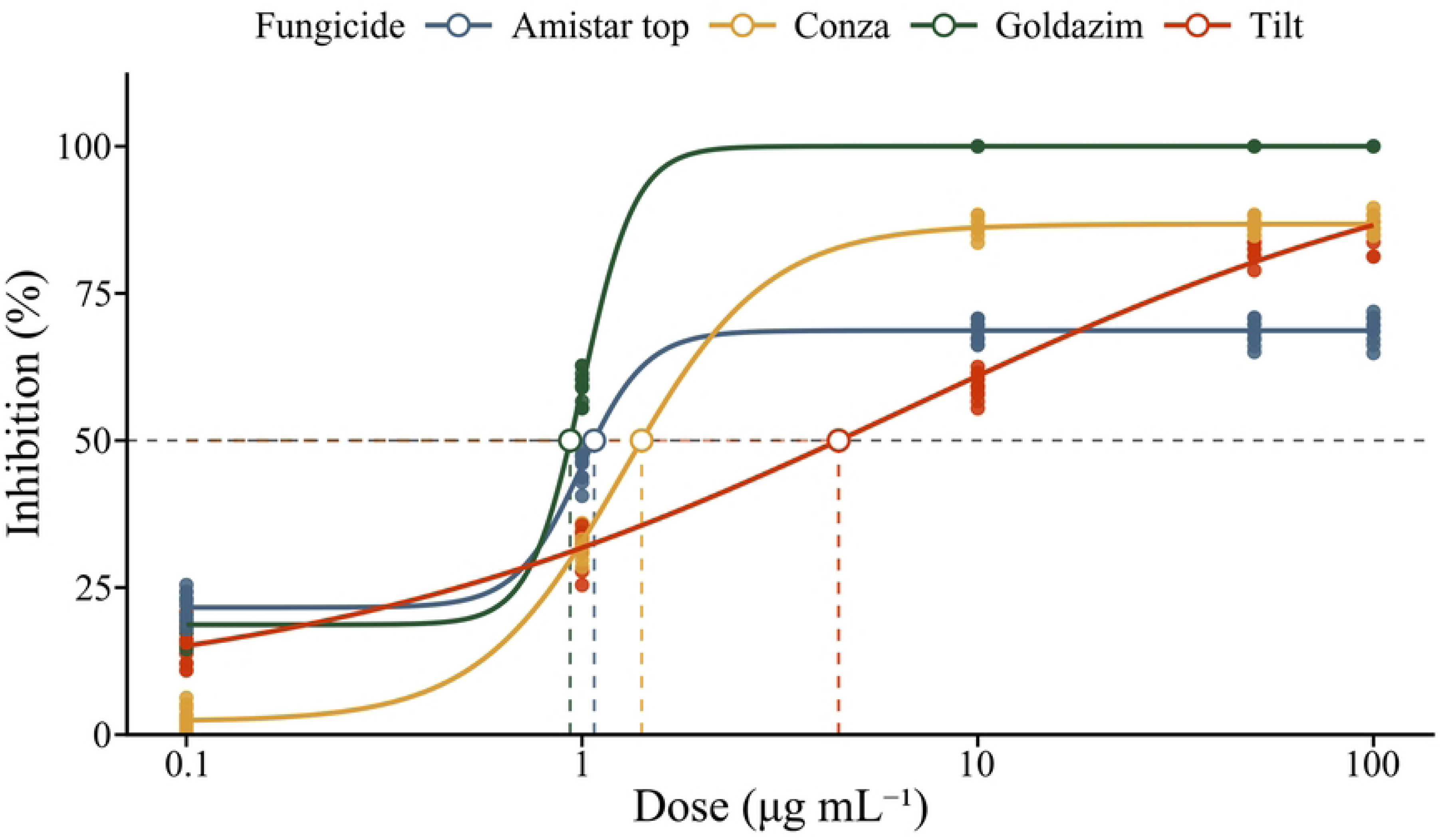
EC_50_(50% mycelial inhibition) determination of fungicides against *C. cassiicola* based on mycelial growth inhibition. Dose-response curves show the relationship between fungicide concentration (μg mL⁻¹) and mycelial growth inhibition for Amistar Top, Conza, Goldazim, and Tilt fungicides. Solid lines represent fitted models for each fungicide. Dots represent observed inhibition values from individual replicates. Open circles indicate the estimated EC_50_ for each fungicide, and dashed lines denote the corresponding 50% inhibition level and EC_50_ positions.

## Discussion

This study for the first time demonstrated the occurrence of target spot disease on chia caused by *C. cassiicola* in Bangladesh. The disease symptoms observed in the present study were typical of Corynespora target spot reported in many other crops [13,34]. The emergence of this pathogen on chia is of particular significance because chia is gaining importance as a high-value food crop in the region. Apart from Bangladesh, this disease is only reported on chia in India, indicating that the pathogen may be expanding its host range and geographical distribution in South Asia [34]. In Bangladesh, *C. cassiicola* has mainly been associated with diseases of okra [35], and its occurrence on chia suggests a potential host shift or adaptation that could pose new risks to crop production.

Morphological characterization of the pathogen supported preliminary its identification as *Corynespora sp*. The isolates in this study produced conidia of variable size and shape, occasionally forming short chains and containing multiple pseudosepta, most commonly 2–5, which is consistent with the known morphological features of the genus *Corynespora* [15,32,36]. However, a predominance of cylindrical and straight conidia was observed in our study, similar to those previously described for this pathogen isolated from soybean [13], whereas some other studies have reported predominantly obclavate and curved conidia in culture media [16,37]. However, the uniformity of conidial morphology among the isolates suggests that a single species was responsible for the disease outbreak on chia in the study area.

Molecular characterization along with morphological characterization is more reliable for fungal species identification which exhibits cryptic morphology [38]. The internal transcribed spacer (ITS) region and the translation elongation factor 1-alpha *(TEF-1α)* gene are widely used as molecular barcodes for fungal species identification [34,39,40]. Although the ITS region is effective for genus-level identification, it may lack sufficient variability to distinguish closely related species within certain genera [41,42]. Therefore, sequencing two or more loci provides more reliable and comprehensive species identification. The *TEF-1α* gene has high phylogenetic resolution and is particularly useful for differentiating cryptic species within a genus [43,44]. Phylogenetic analysis based on combined ITS and *TEF-1α* sequences of two representative isolates showed that the isolates clustered with reference strains of *C. cassiicola*. This multilocus approach provided strong molecular evidence and further confirmed the identity of the pathogen.

Pathogenicity tests were performed to determine the virulence of the *C. cassiicola* isolates, and the results clearly demonstrated their ability to infect chia and reproduce the characteristic target spot symptoms observed under field conditions. The inoculated plants developed typical concentric necrotic lesions, thereby fulfilling Koch’s postulates and confirming the causal role of the pathogen. *C. cassiicola* is widely recognized as a destructive and highly adaptable pathogen, reported to cause leaf spot, defoliation, and inflorescence blight in more than 500 plant species worldwide [12]. Its broad host range and aggressive infection strategy make it particularly concerning in diversified cropping systems.

In the present study, *C. cassiicola* was identified for the first time as the causal agent of target spot disease of chia in Bangladesh, highlighting the emergence of a new host–pathogen association in the country. More alarmingly, cross-pathogenicity assays revealed that the isolates were capable of infecting other economically important vegetable crops cultivated in Bangladesh (Fig 5). This finding suggests that the pathogen possesses a flexible host adaptation capacity, increasing the risk of spillover to additional crops. Given Bangladesh’s intensive and often overlapping vegetable production systems, such cross-infectivity may facilitate rapid dissemination across fields and seasons [45]. If not properly managed, the pathogen could establish itself in multiple hosts, creating a persistent inoculum reservoir and escalating disease pressure. Therefore, the emergence of *C. cassiicola* on chia should be viewed not as an isolated incident but as a potential threat to a wider range of vegetable crops in Bangladesh, warranting immediate surveillance and integrated disease management strategies to prevent further spread and host expansion.

The management of target spot disease largely relies on chemical control, particularly under environmental conditions that favor rapid disease development. In the present study, seven fungicides with different modes of action were evaluated in vitro to determine their effectiveness against the pathogen. Among them, Goldazim had a lower EC_50_ value (0.76 μg mL⁻¹) and complete suppression was achieved at 10 μg mL⁻¹), indicating high antifungal activity. The superior performance of the fungicide suggests that this could be promising candidate for the management of target spot disease of chia under field conditions. Goldazim contains Carbendazim, a systemic benzimidazole fungicide, known for its effectiveness against a wide range of fungal pathogens, including *C. cassiicola*. This fungicide interferes with fungal cell division by disrupting the polymerization of monomeric tubulins, thereby inhibiting mitosis, which likely contributed to its high efficacy [46]. However, despite its strong antifungal activity, the development of resistance to carbendazim in *C. cassiicola* populations has been reported [18].

The response of *C. cassiicola* to the Conza and Amistar Top indicates a clear distinction between potency and maximum efficacy. The mixture of azoxystrobin + difenoconazole (Amistar Top) showed a lower EC₅₀, suggesting greater activity at low concentrations, but resulted in lower inhibition at 10 µg ml⁻¹ compared with hexaconazole (Conza). This suggests that the mixture is more potent but has a limited maximum inhibitory effect. In contrast, hexaconazole achieved greater inhibition at higher concentration (10 µg ml⁻¹), indicating stronger overall efficacy. These differences may be attributed to their distinct modes of action. Azoxystrobin, a QoI fungicide, inhibits mitochondrial respiration by blocking electron transport, while difenoconazole and hexaconazole, both triazoles, interfere with ergosterol biosynthesis [47–49]. The combined action in the mixture of azoxystrobin + difenoconazole may result in strong initial growth suppression but could be constrained by partial resistance, metabolic adaptation, or reduced sensitivity of *C. cassicola*, thereby limiting its maximal effect. Conversely, hexaconazole may exert a more sustained inhibitory effect on membrane integrity at higher concentrations, leading to greater overall growth suppression. Interestingly, only a few studies have evaluated the hexaconazole fungicides against *C. cassiicola*, and those studies reported effective disease control under both in vitro and field conditions [50,51].Additionally, differences in fungistatic versus fungicidal activity, uptake efficiency, and detoxification mechanisms in the pathogen may further contribute to the observed response. Therefore, EC₅₀ values alone may not fully represent fungicide performance against *C. cassicola*.

However, it is important to note that reliance solely on fungicides may not be sustainable in the long term due to the risk of fungicide resistance development in *C. cassiicola*. This pathogen highly adaptable and genetically diverse and is classified by FRAC as “high risk” for developing fungicide resistance [12]. Therefore, integrated disease management strategies, including crop sanitation, use of disease-free planting material, monitoring of environmental conditions, and rational fungicide rotation, should be considered to minimize disease pressure and prolong fungicide efficacy. Further field-based evaluations are necessary to validate the in vitro findings and to optimize application timing and dosage for effective disease control.

## Conclusion

This study reports, for the first time, the occurrence of *C. cassiicola*, the causal agent of target spot disease in chia, in Bangladesh. The findings highlight the importance of continuous surveillance of emerging diseases in newly introduced or expanding crops. Monitoring the host range of the pathogen through field surveys will also be important for designing effective crop rotation strategies and crop isolation practices to limit disease spread. The fungicide sensitivity assay further demonstrated the effectiveness of three fungicides Goldazim, Conza and Amistar Top in suppressing the growth of *C. cassiicola*. Given the increasing cultivation and economic importance of chia, early detection and effective management of target spot disease will be critical to prevent potential yield losses and ensure sustainable crop production. Future research focusing on pathogen diversity, epidemiology, fungicide resistance, and integrated disease management strategies will further improve our understanding and help develop effective control measures for this emerging disease.

## Author Contributions

Avi Kumar Badhon: investigation, methodology, formal analysis, writing – original draft. Dipali Rani Gupta: methodology, validation, supervision, writing – review and editing. Swapan Kumar Paul: sampling, data curation, validation, writing – review and editing. Julfikar Ali methodology, data curation. Md. Moshiur Rahman: sampling, writing – review and editing. Tofazzal Islam: conceptualization, validation, supervision, resource, writing – review and editing.

## Data Availability Statement

The newly obtained sequences of the internal transcribed spacer (ITS) regions (including 1 and 2 and intervening 5.8S nrDNA) and the translation elongation factor 1- α (*tef1*) gene have been deposited in the NCBI GenBank database.

## References

1. Shorna SI, Akhter BA, Anwar MP, Hossain MA. Growth and yield response of BAU Chia-1 (*Salvia hispanica*) to nitrogen fertilizer. J Bangladesh Agric Univ. 2024 Dec 31;22(4):443–50. doi:10.3329/jbau.v22i4.78849

2. Khalid W, Arshad MS, Aziz A, Rahim MA, Qaisrani TB, Afzal F, et al. Chia seeds (*Salvia hispanica* L.): A therapeutic weapon in metabolic disorders. Food Sci Nutr. 2023 Jan;11(1):3–16. doi:10.1002/fsn3.3035

3. Motyka S, Skała E, Ekiert H, Szopa A. Health-promoting approaches of the use of chia seeds. J Funct Foods. 2023 Apr;103:105480. doi:10.1016/j.jff.2023.105480

4. Anand G, Rajeshkumar KC. Challenges and threats posed by plant pathogenic fungi on agricultural productivity and economy. In: Rajpal VR, Singh I, Navi SS, editors. Fungal diversity, ecology and control management. Singapore: Springer Nature Singapore; 2022. pp. 483–93. https://link.springer.com/10.1007/978-981-16-8877-5_23

5. Gai Y, Wang H. Plant disease: A growing threat to global food security. Agronomy. 2024 Jul 24;14(8):1615. doi:10.3390/agronomy14081615

6. Bebber DP, Ramotowski MAT, Gurr SJ. Crop pests and pathogens move polewards in a warming world. Nat Clim Change. 2013 Nov;3(11):985–8. doi:10.1038/nclimate1990

7. Karunarathna SC, Maharachchikumbura SSN, Ariyawansa HA, Shenoy BD, Jeewon R. Editorial: Emerging fungal plant pathogens. Front cell infect microbiol. 2021 Sep;11:765549. doi:10.3389/fcimb.2021.765549

8. Singh BK, Delgado-Baquerizo M, Egidi E, Guirado E, Leach JE, Liu H, et al. Climate change impacts on plant pathogens, food security and paths forward. Nat Rev Microbiol. 2023 Oct;21(10):640–56. doi:10.1038/s41579-023-00900-7

9. Gange AC, Gange EG, Mohammad AB, Boddy L. Host shifts in fungi caused by climate change? Fungal Ecol. 2011 Apr;4(2):184–90. doi:10.1016/j.funeco.2010.09.004

10. Termorshuizen AJ. Ecology of fungal plant pathogens. Microbiol Spectr. 2016 Dec;4(6):10–128. doi:10.1128/microbiolspec.FUNK-0013-2016

11. Islam MT, Croll D, Gladieux P, Soanes DM, Persoons A, Bhattacharjee P, et al. Emergence of wheat blast in Bangladesh was caused by a South American lineage of *Magnaporthe oryzae*. BMC Biol. 2016 Dec;14(1):84. doi:10.1186/s12915-016-0309-7

12. N. Rondon M, Lawrence K. The fungal pathogen *Corynespora cassiicola*: A review and insights for target spot management on cotton and Soya bean. J Phytopathol. 2021 Jun;169(6):329–38. doi:10.1111/jph.12992

13. Toulet ML, Neira DA, Escobar M, Pardo EM, Arias ME, Ploper LD, et al. Morphological and pathogenic characterization of *Corynespora cassiicola* isolates reveals specific genotypic interactions in soybean. Plant Pathol. 2022 May;71(4):843–59. doi:10.1111/ppa.13528

14. Feng R, Wang H, Zhang X, Li T, Huang C, Zhang S, et al. Characteristics of *Corynespora cassiicola*, the causal agent of tobacco Corynespora leaf spot, revealed by genomic and metabolic phenomic analysis. Sci Rep. 2024 Aug;14(1):18326. doi:10.1038/s41598-024-67510-y

15. Qi YX, Zhang X, Pu JJ, Liu XM, Lu Y, Zhang H, et al. Morphological and molecular analysis of genetic variability within isolates of *corynespora cassiicola* from different hosts. Eur J Plant Pathol. 2011 May;130(1):83–95. doi:10.1007/s10658-010-9734-6

16. Khadka RB, Bhandari R, Rimal A, Gaire A, Adhikari S, Bhandari S, et al. First report of target spot on tomato caused by *Corynespora cassiicola* in Nepal. New Dis Rep. 2023 Jul;48(1):e12187. doi:10.1002/ndr2.12187

17. Zhu F, Shi Y, Xie X, Chai A, Li B. Occurrence, Distribution, and Characteristics of Boscalid-Resistant *Corynespora cassiicola* in China. Plant Dis. 2019 Jan;103(1):69–76. doi:10.1094/PDIS-11-17-1760-RE

18. Deng Y, Wang T, Zhao P, Du Y, Zhang L, Qi Z, et al. Sensitivity to 12 fungicides and resistance mechanism to trifloxystrobin, carbendazim, and succinate dehydrogenase inhibitors in cucumber corynespora leaf spot (*Corynespora cassiicola*). Plant Dis. 2023 Dec;107(12):3783–91. doi:10.1094/PDIS-04-23-0615-RE

19. Sun B, Zhou R, Zhu G, Xie X, Chai A, Li L, et al. The mechanisms of target and non-target resistance to QoIs in *Corynespora Cassiicola*. Pestic Biochem Physiol. 2024 Jan;198:105760. doi:10.1016/j.pestbp.2023.105760

20. Duan Y, Xin W, Lu F, Li T, Li M, Wu J, et al. Benzimidazole- and QoI-resistance in *Corynespora cassiicola* populations from greenhouse-cultivated cucumber: An emerging problem in China. Pestic Biochem Physiol. 2019 Jan;153:95–105. doi:10.1016/j.pestbp.2018.11.006

21. Miyamoto T, Ishii H, Seko T, Kobori S, Tomita Y. Occurrence of *Corynespora cassiicola* isolates resistant to boscalid on cucumber in Ibaraki Prefecture, Japan. Plant Pathol. 2009 Dec;58(6):1144–51. doi:10.1111/j.1365-3059.2009.02151.x

22. Zhao Q, Shi Y, Wang Y, Xie X, Li L, Fan T, et al. Temperature and humidity regulate sporulation of *Corynespora cassiicola* that is associated with pathogenicity in cucumber (*Cucumis sativus* L.). Biology. 2022 Nov;11(11):1675. doi:10.3390/biology11111675

23. White TJ, Bruns T, Lee S, Taylor J. Amplification and direct sequencing of fungal ribosomal rna genes for phylogenetics. In: Innis MA, Gelfand DH, Sninsky JJ, White TJ, editors. PCR Protocols: a guide to methods and applications. 1990. pp. 315–22. doi:10.1016/B978-0-12-372180-8.50042-1

24. Surovy MZ, Kabir MK, Gupta DR, Hassan O, Mahmud NU, Sabir AA, et al. First report of fusarium wilt caused by *Fusarium oxysporum* on strawberry in Bangladesh. Plant Dis. 2019 Feb;103(2):367. doi:10.1094/PDIS-07-18-1121-PDN

25. Carbone I, Kohn LM. A method for designing primer sets for speciation studies in filamentous ascomycetes. Mycologia. 1999 May;91(3):553–6. doi:10.1080/00275514.1999.12061051

26. Katoh K, Standley DM. MAFFT multiple sequence alignment software version 7: improvements in performance and usability. Mol Biol Evol. 2013 Apr;30(4):772–80. doi:10.1093/molbev/mst010

27. Tamura K, Stecher G, Kumar S. MEGA11: molecular evolutionary genetics analysis version 11. Mol Biol Evol. 2021 Jun;38(7):3022–7. doi:10.1093/molbev/msab120

28. Zhang C, Imran M, Xiao L, Hu Z, Li G, Zhang F, et al. Difenoconazole resistance shift in *Botrytis cinerea* from tomato in China associated with inducible expression of *CYP51*. Plant Dis. 2021 Feb;105(2):400–7. doi:10.1094/PDIS-03-20-0508-RE

29. Cerritos-Garcia DG, Huang SY, Kleczewski NM, Mideros SX. Virulence, aggressiveness, and fungicide sensitivity of *Phytophthora* spp. associated with soybean in Illinois. Plant Dis. 2023 Jun ;107(6):1785–93. doi:10.1094/PDIS-07-22-1551-RE

30. Song Y, Li L, Li C, Lu Z, Men X, Chen F. Evaluating the sensitivity and efficacy of fungicides with different modes of action against *Botryosphaeria dothidea*. Plant Dis. 2018 Sep;102(9):1785–93. doi:10.1094/PDIS-01-18-0118-RE

31. Noel ZA, Wang J, Chilvers MI. Significant Influence of EC_50_ estimation by model choice and EC_50_ type. Plant Dis. 2018 Apr;102(4):708–14. doi:10.1094/PDIS-06-17-0873-SR

32. Ellis MB, Holliday P. Corynespora cassiicola. Kew, UK: Commonwealth Mycological Institute; 303 pp [Internet]. 1971.

33. Manjunatha N, Pokhare SS, Sharma J, Patil PG, Agarrwal R, Chakranarayan MG, et al. Collar rot caused by *Calonectria hawksworthii*, a new record for pomegranate (*Punica granatum*). J Plant Pathol. 2023 May;105(3):887–94. doi:10.1007/s42161-023-01391-4

34. Manjunatha H, Govind G, Channakeshava C, Shashi Kiran AS, Koler P. First evidence of *Corynespora cassiicola*-induced disease in Chia: A new challenge for Indian Chia cultivation. Physiol Mol Plant Pathol. 2025 Nov;140:102917. doi:10.1016/j.pmpp.2025.102917

35. Ahmed FA, Alam N, Khair A. Incidence and biology of *Corynespora cassiicola* (Berk. & Curt.) Wei. disease of okra in Bangladesh. Bangladesh J Bot. 2014 Feb;42(2):265–72. doi:10.3329/bjb.v42i2.18028

36. Xu J, Gong G, Cui Y, Zhu Y, Wang J, Yao K, et al. Comparison and correlation of *Corynespora cassiicola* populations from Kiwifruit and other hosts based on morphology, phylogeny, and pathogenicity. Plant Dis. 2023 Jul;107(7):1979–92. doi:10.1094/PDIS-04-22-0937-RE

37. Sain SK, Gawande SP, Kumar V, Chandrashekar N, Prakash AH, Prasad YG. First report of target spot caused by *Corynespora cassiicola* on Cotton in Northwestern India. Plant Dis. 2024 Feb;108(2):530. doi:10.1094/PDIS-09-23-1868-PDN

38. Ekanayaka AH, Karunarathna SC, Tibpromma S, Dutta AK, Tennakoon DS, Karunarathna A, et al. Species evolution: cryptic species and phenotypic noise with a particular focus on fungal systematics. Front Cell Infect Microbiol. 2025 Feb;15:1497085. doi:10.3389/fcimb.2025.1497085

39. Wang Y, Hao D, Jiang H, Fei Z, Zhao R, Gao J, et al. Identification and fungicide sensitivity of *Fusarium oxysporum*, the cause of Fusarium wilt on *Codonopsis pilosula* in Shanxi Province (Lu Dangshen), China. Crop Prot. 2025 Apr;190:107083. doi:10.1016/j.cropro.2024.107083

40. Kashyap PL, Rai P, Kumar S, Chakdar H, Srivastava AK. Erratum to: DNA barcoding for diagnosis and monitoring of fungal plant pathogens. In: Singh BP, Gupta VK, editors. Molecular Markers in Mycology. Cham: Springer International Publishing; 2017. pp. E1–E1. doi:10.1007/978-3-319-34106-4_17

41. Karlsson I, Edel-Hermann V, Gautheron N, Durling MB, Kolseth AK, Steinberg C, et al. Genus-specific primers for study of *Fusarium* communities in field samples. Appl Environ Microbiol. 2016 Jan;82(2):491–501. doi:10.1128/AEM.02748-15

42. Wang Y, Chen JY, Xu X, Cheng J, Zheng L, Huang J, et al. Identification and characterization of *Colletotrichum* species associated with anthracnose disease of *Camellia oleifera* in China. Plant Dis. 2020 Feb;104(2):474–82. doi:10.1094/PDIS-11-18-1955-RE

43. Boutigny AL, Gautier A, Basler R, Dauthieux F, Leite S, Valade R, et al. Metabarcoding targeting the EF1 alpha region to assess *Fusarium* diversity on cereals. PLOS ONE. 2019 Jan;14(1):e0207988. doi:10.1371/journal.pone.0207988

44. O’Donnell K, Kistler HC, Tacke BK, Casper HH. Gene genealogies reveal global phylogeographic structure and reproductive isolation among lineages of *Fusarium graminearum*, the fungus causing wheat scab. Proc Natl Acad Sci. 2000 Jul;97(14):7905–10. doi:10.1073/pnas.130193297

45. Gupta R, Sarkar MJ, Islam MdS, Uddin MdR, Riza IJ, Monira S, et al. Introducing New Cropping Pattern to Increase Cropping Intensity in Hill Tract Area in Bangladesh. Sustainability. 2023 Jul;15(14):11471. doi:10.3390/su151411471

46. Zhou F, Zhou X, Jiao Y, Han A, Su H, Wang L, et al. Potential mechanisms of hexaconazole resistance in *Fusarium graminearum*. Plant Dis. 2024 Oct;108(10):3133–45. doi:10.1094/PDIS-04-24-0880-RE

47. Zhou Y, Xu J, Zhu Y, Duan Y, Zhou M. Mechanism of action of the benzimidazole fungicide on *Fusarium graminearum*: Interfering with polymerization of monomeric tubulin but not polymerized microtubule. Phytopathology. 2016 Aug;106(8):807–13. doi:10.1094/PHYTO-08-15-0186-R

48. Liu Y, Tan X, Zhao J, Niu Y, Hsiang T, Yu Z, et al. Diversity of *Colletotrichum* species associated with anthracnose on *Euonymus japonicus* and their sensitivity to fungicides. Front Plant Sci. 2024 Jun;15:1411625. doi:10.3389/fpls.2024.1411625

49. Ghannoum MA, Rice LB. Antifungal agents: mode of action, mechanisms of resistance, and correlation of these mechanisms with bacterial resistance. Clin Microbiol Rev. 1999 Oct;12(4):501–17. doi:10.1128/CMR.12.4.501

50. Kurre AK. Studies on variability and management of target leaf spot of soybean caused by Corynespora cassiicola (Berk. and Curt.) Wei. M. Sc.(Agri.) Thesis, Indira Gandhi Krishi Vishwavidyalaya; 2016. Available from: https://krishikosh.egranth.ac.in/server/api/core/bitstreams/deef3555-ef21-4be4-b07a-f09022e529c4/content

51. Molina JPE. Yield losses of soybean due to target spot (Corynespora cassiicola), its genetic and chemical management. Doctoral dissertation, Universidade de São Paulo; 2018. Available from: http://www.teses.usp.br/teses/disponiveis/11/11135/tde-25072018-165739/

